# Engineered yeast biofilms deliver plastic degrading enzymes to PET substrates

**DOI:** 10.1101/2025.05.27.655536

**Authors:** Charlotte Rose Bilsby, Samantha K. Hobbs, Jack Davis, Samuel Merryn Jones, Philippe Pierre Laissue, Rosie Graham, Ed Deshmukh-Reeves, Sarina Sanami, Noor Issa, Wei-Feng Xue, Andrew Lawrence, Campbell Gourlay, Andrew R. Pickford, Tobias von der Haar

**Affiliations:** School of Biosciences, Division of Natural Sciences, University of Kent, Canterbury CT2 7NJ, UK; Centre for Enzyme Innovation, School of Biological Sciences, University of Portsmouth, Portsmouth PO1 2DT, UK; School of Life Sciences, University of Essex, Colchester CO4 3SQ, UK; School of Biological Sciences, University of Southampton, Southampton, SO17 1BJ, UK

## Abstract

Enzymes that degrade PET and other plastics are being discovered with increasing pace, and engineering approaches further increase their activity. Most proposed applications for such enzymes require their purification, enabling the regeneration of monomeric subunits suitable for the production of new plastic. However, large amounts of plastics exist as environmental contaminants, and are unlikely to be suitable for standard recycling. Such waste may still be suitable for bioremediation using enzymatic activities, albeit without the economic incentives of a full recycling process. Here we develop a delivery system for PET hydrolases, based on engineered yeast biofilms that grow and secrete enzymes directly on the substrate to be degraded. The system does not require any enzyme purification steps, and the biofilms at least partially assimilate the degradation products into biomass. Our work provides proof-of-principle that demonstrates the potential of biofilm-based approaches.

## Introduction

Plastics include some of the most widely used synthetic materials and constitute key environmental pollutants. A high proportion of plastics are produced for single use applications, including polyethylene terephthalate (PET)-based beverage and food containers. Although rates of recycling of PET-based materials are increasing, historical recycling rates are low, and only 9% of all plastics ever produced have been estimated to be recycled into secondary materials or products (1). The remainder have been discarded to landfill or into the environment.

Once discarded, the lifetime of PET materials is estimated to be on the scale of decades (2) or centuries (3). Degradation of macroscopic items frequently involves release of smaller PET particles in the millimetre to micrometre range (microplastics), a process that is accelerated by agitation for example in marine surf (4) or in areas subject to wind abrasion (5). Following decades of uncontrolled plastic disposal, plastic and microplastic contaminants are now found in all areas of the environment, from remote mountain top lakes (6) to the gastrointestinal tract of benthic deep-sea organisms (7) and inside humans (8). While both technical and consumer behaviour aspects of recycling are intensely researched and this can help to reduce rates of the introduction of new plastic waste into the environment, it does not address existing environmental contamination. Large-scale clean-up operations are underway to recover macroscopic plastic waste from the marine environment, but similar efforts for terrestrial contamination are not yet emerging, and microplastics remain challenging to remove from any environment (9,10).

The discovery of PET hydrolases that naturally degrade PET with moderate activity, such as *Ideonella sakaiensis* PETase (*Is*PETase) and Leaf Compost Cutinase (LCC) (11–14), coupled with the application of engineering approaches to increase the specific activity of these enzymes (15,16), could be a promising solution for the bioremediation of environmental plastic contaminants.

Since expression levels in their native organisms are too low for industrial applications, a variety of heterologous expression systems have been developed for PET hydrolases, with some representative examples shown in table 1. Some of these systems aim at the high-level production of enzymes for purification, and can now reach gram per litre yields for the most optimised expression systems. Other applications aim to enable PET hydrolase delivery in non-purified form, for example as whole cell catalysts where the enzymes are linked to the cell wall of the expression host (17,18). Such systems typically generate lower expression yields, which are nevertheless compatible with their specific intended purpose.

**Table 1.**
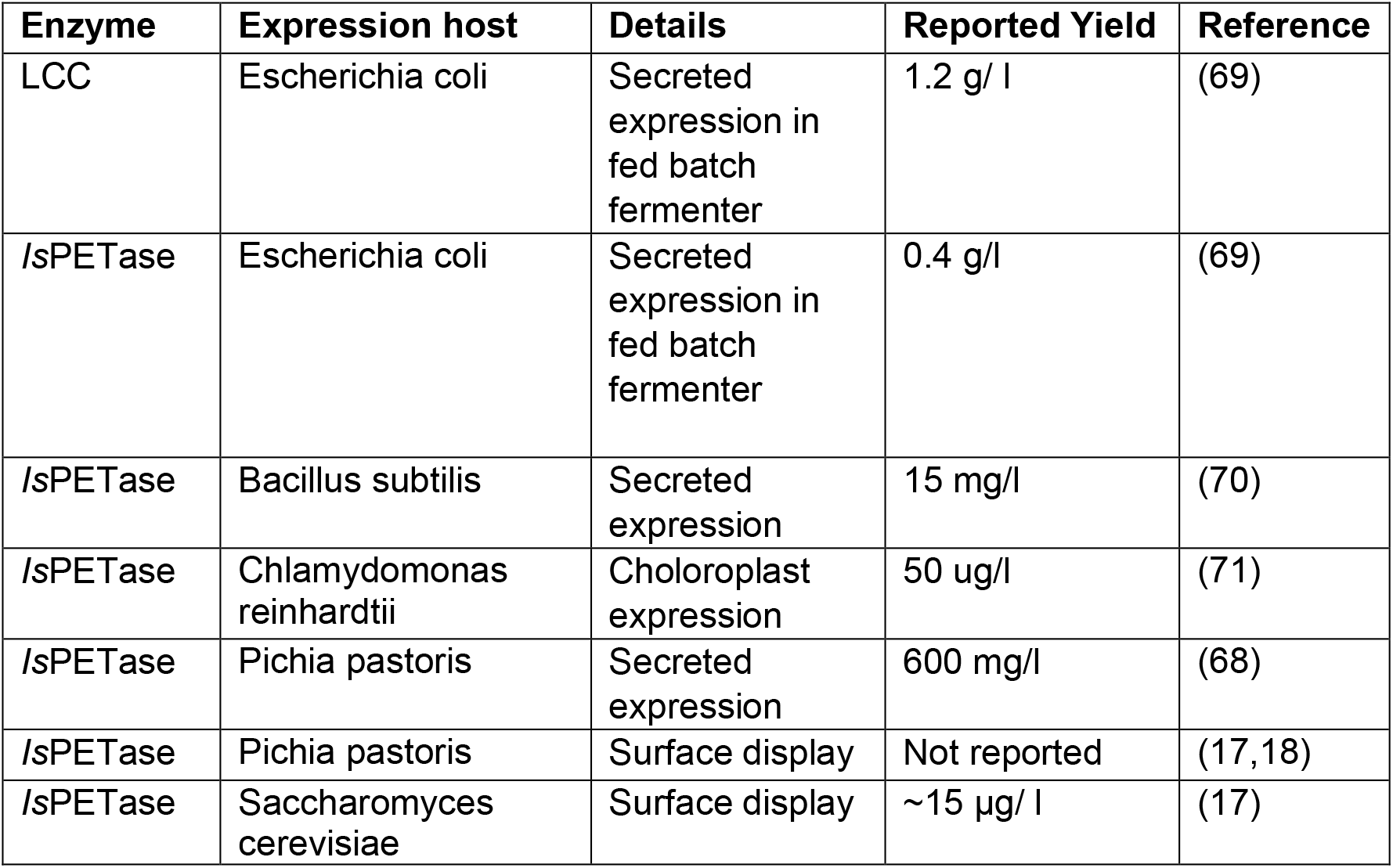
Representative host systems for expression of PET hydrolases.

The most common proposed applications for PET hydrolases involve their purification and application to clean post-consumer plastic, which enables the generation of PET monomers suitable for the re-polymerisation of virgin PET (19). While this approach is promising for clean post-consumer waste, existing environmental plastic contaminants are typically of mixed composition including environmental debris, from which they cannot easily be separated. Moreover, they may be chemically altered through the influence of UV and oxygen (20), which may make them difficult or impossible to fully recycle (21). PET hydrolases could still be used to remove such contaminants and turn them into bio-accessible breakdown products, but if these products are then not used to produce new PET the process would lose financial incentive, and the additional costs of enzyme purification could be a deterrent from applying enzyme-based bioremediation strategies.

For applications where the main intention is bioremediation, for example the enzymatic removal of contaminated plastic recovered from the environment, enzyme purification would in principle not be necessary if the enzymes were active in crude cell extracts or growth supernatants. However, achieving the high enzyme-to-plastic ratios required for efficient degradation (optimally above 2 mg enzyme per g plastic (22)), would be challenging in such formats. We reasoned that one way of addressing this limitation could be through the formation of biofilms by the expression host on the degradation substrate.

Biofilms are formed by the attachment of cells to surfaces and their concomitant growth and embedding into extracellular matrices (23). They can be formed by single species or can be polymicrobial (24), and are prevalent in a diverse range of ecosystems, spanning both marine (25) and terrestrial environments (26). Biofilm-embedded cells can often withstand environmental and other stresses better than planktonic cells, including sub-optimal pH and salinity (27). Moreover, growth in biofilms can improve productivity in recombinant protein expression systems (28,29). Surface hydrophobicity is a dominant characteristic determining the efficiency of biofilm formation (30,31), meaning that plastic materials are naturally good biofilm substrates. Indeed, environmental plastic is frequently colonised by both microscopic and macroscopic organisms, giving rise to the notion of the “plastisphere” (32).

Yeasts and other fungi are particularly active in biofilm formation (33) as well as having well defined pathways for secreting proteins into extracellular space (34). Their biofilm matrix is composed of extracellular polymeric substances (EPS), secreted proteins and other biomolecules that embed the cells and stabilise the biofilm (35). Diffusion within the biofilm matrix is significantly more limited compared to liquid medium (36), and one can thus make the rational argument that the secretion of plastic degrading enzymes within a yeast biofilm matrix should lead to an increase in enzyme concentration close to the plastic surface. Recent results on the association rate between *Is*PEtase and PET surfaces, which reported results as low as 1000 association events per second per μm^2^ and M enzyme (37), indicated that increasing enzyme concentration close to the plastic surface could be a particularly powerful way of enhancing degradation rates.

With these considerations in mind, we developed yeast secretion systems that deliver active PET hydrolases into the growth medium. We observed evidence of plastic degradation both with planktonic cells and with biofilms, with significantly stronger degradation activity observed with biofilms. Overall, our work provides proof of principle for the application of engineered biofilms in the remediation of environmental plastic contaminants and enable us to define requirements for the development of biofilm-based systems with sufficiently high activity levels to enable actual environmental applications.

## Results

### Expression of secreted IsPETase in baker’s yeast

To explore whether baker’s yeast was able to secrete PET hydrolases, we initially introduced expression optimised genes for *IsPETase* into two common laboratory yeast strains, BY4741 and Σ1278b. Following growth of the transformed strains in defined medium we tested for anti-HA reactive bands in both cell extracts and growth supernatants (Figure 1). The theoretical molecular weights are 36.2 kDa for the full-length protein and 31.4 kDa following cleavage of the signal sequence. As seen in figure 1B, an HA-reactive band around the correct molecular weight for the latter product was observed in cell extracts. In addition, we also observed higher molecular weight bands in cell extracts, and supernatant samples exclusively showed higher molecular weight bands.

**Figure 1.**
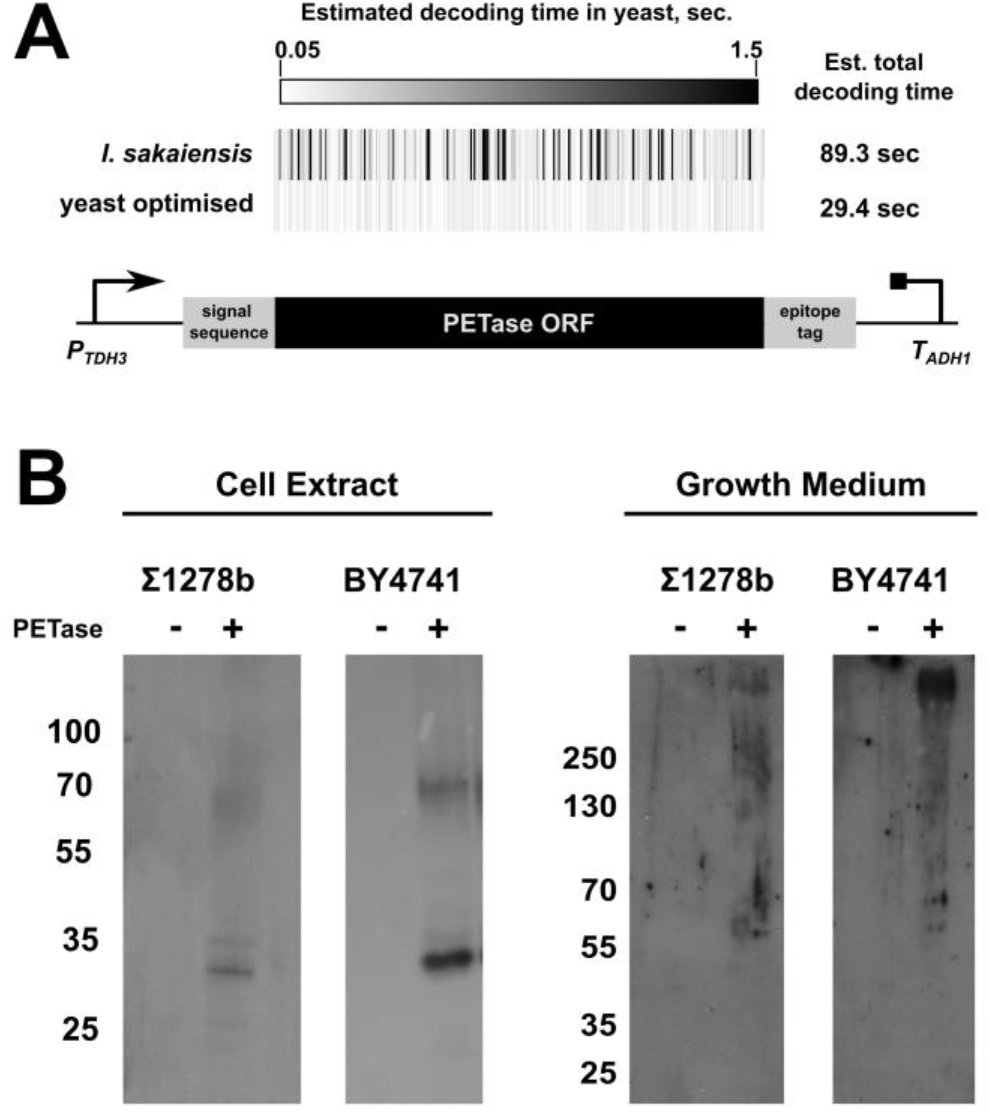
*Is*PETase expression in baker’s yeast. **A**, codon optimality and predicted decoding times in yeast for the native *I. sakaiensis* PET hydrolase gene and the yeast optimised gene version used here. **B**, anti-HA western blots of cell extracts (left) and supernatant samples (right) of two yeast strains. Numbers indicate position of molecular weight markers (kDa). +, PET hydrolase vector transformants; -, empty vector transformants.

Baker’s yeast modifies some secreted proteins through addition of O- and N-linked glycans (47), and we reasoned that the higher molecular weight bands in the supernatant could correspond to glycosylated *Is*PETase. Consistent with this hypothesis, a computational glycosylation prediction tool (48) identified 5 asparagines in the *Is*PETase sequence as high confidence glycosylation substrates (quality scores >0.9, Figure 2). To further test whether the high molecular weight bands corresponded to glycosylated *Is*PETase, protein samples precipitated from the growth supernatant were treated with Endo Hf, an enzyme that cleaves asparagine-linked high-mannose glycans (49). Following Endo Hf treatment, a significant reduction in molecular weight was observed, with a distinct central band migrating around the molecular weight of the main intracellular product (Figure 2). We conclude that the high molecular weight bands observed in the cell culture supernatants correspond to glycosylated *Is*PETase.

**Figure 2.**
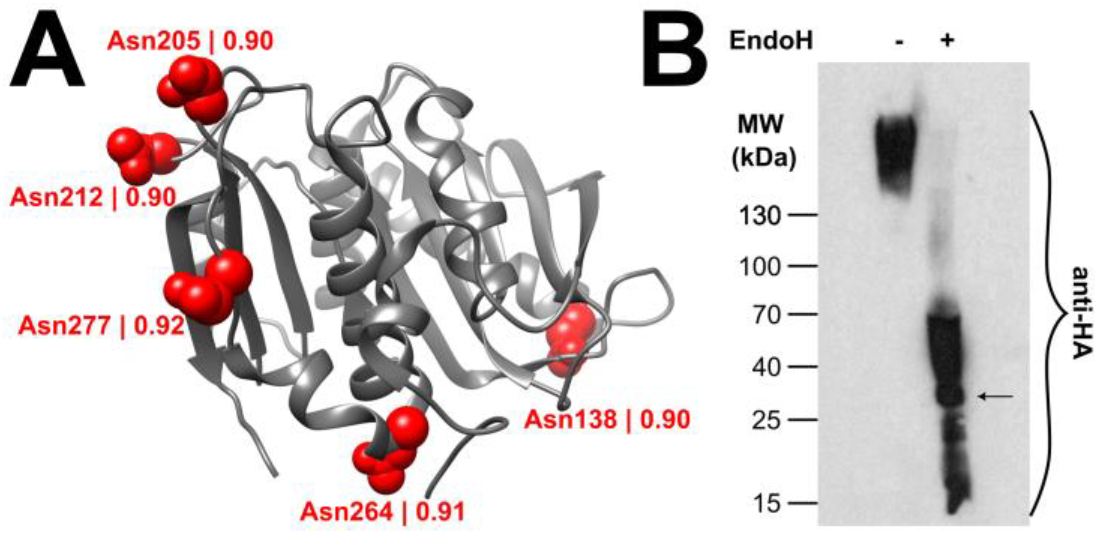
Secreted PET hydrolase is glycosylated. **A**, High-confidence glycosylation sites in *I. sakaiensis* PET hydrolase predicted using MusiteDeep. **B**, anti-HA western blots of secreted PET hydrolase samples. -, untreated sample; +, sample treated with recombinant Endoglycosidase H.

Glycosylation is frequently introduced into proteins in response to incomplete folding (50). The modification increases the solubility of its targets and enhances their ability to fold correctly (51). *Is*PETase has been engineered for higher thermostability, mostly with the aim of enabling degradation above the glass transition temperature of PET, where PET polymer chains can be more easily accessed by the enzyme. In addition to allowing enzyme activity at higher temperatures, higher thermostability can also result in more stable conformations at intermediate temperatures. We therefore explored whether the degree of glycosylation in yeast *in vivo* was altered for PET hydrolase variants that had been selected for increased thermostability. Indeed, when we compared levels of secretion of wild type *Is*PETase (T_m_ 45°C), *Is*PETase^Ts^ (T_m_ 56.8°C)(52) and hotPETase (T_m_ 80.5°C)(53) we observed a gradual increase in expression coincident with a higher proportion of less extensively glycosylated protein. This effect was particularly pronounced for the hotPETase variant (figure 3). In addition, two variants of another PET hydrolase, the generally very thermostable Leaf Compost Cutinase (LCC, T_m_ 86°C (54); and LCC^ICCG^, T_m_ 91.7°C (53)), were both expressed without any apparent glycosylation, with the more stable variant showing higher secretion levels (figure 3). These results confirm the connection between folding stability and secretion. Interestingly, expression of LCC in *Pichia pastoris* was shown to produce glycosylated enzyme (54) whereas we observed no glycosylation in *S. cerevisiae*, indicating that the presence or absence of glycosylation is also organism- and context-specific.

**Figure 3.**
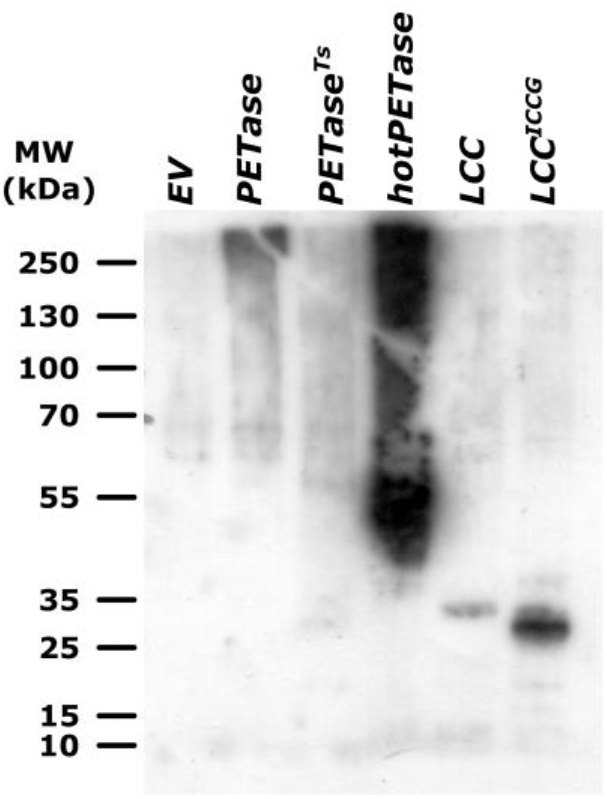
Secreted expression yields of different PET hydrolases. Engineered variants with differing thermostability of *Is*PETase and Leaf Compost Cutinase (LCC) were expressed in baker’s yeast. Expression levels were analysed in the growth supernatant following over night growth.

We estimated secretion yields for HA-tagged hotPETase, the most highly expressed of our PET hydrolases, by comparing western blot signals of growth supernatants from a hotPETase expressing strain against HA-tagged protein standards of know concentration (supplemental figure 1). From these data we estimate that hotPETase is secreted to levels of 400 μg/l. In comparison to other expression systems these levels are at the lower end (table 1), but they are comparable with other specialised expression systems where the primary aim is to enable particular deployment modes for the enzyme, rather than bulk enzyme production for purification. If biofilms were able to strongly concentrate the secreted enzyme in the vicinity of the degradation target, these expression levels should be sufficient to achieve PET degradation provided the secreted enzymes are active.

### Yeast biofilm formation on PET substrates

We investigated the ability of two yeast strains to form biofilms on PET surfaces, of which one (Σ1278b) has been reported to form biofilms efficiently (55), whereas the other (BY4741) does not. Post-consumer PET samples were incubated with logarithmically growing cultures of these two strains, and cell adherence to the plastic surface quantified via a crystal violet staining assay. In this assay Σ1278b attached to the PET surface more efficiently than BY4741 (figure 4A). We investigated the nature of this attachment in more detail by examining early (1 day-old) biofilms of Σ1278b grown on PET coupons, where yeast cells were stained with calcofluor white (CFW). At this early-stage yeast form small biofilm seeds distributed over the entire plastic surface (figure 4B). The samples had been washed with buffer prior to imaging, demonstrating that attachment of these seed biofilms to the surface is stable. Comparison of colony sizes on the smooth coupon surface with colonies on the rough outer edge where the coupon had been cut revealed that seeds were significantly larger in the latter area (figure 4C, average diameter 4.7 μm in the rough area *vs*. 2.3 μm in smooth), indicating that seeds form more easily around crevices and other surface faults. Moreover, when we investigated PET samples incubated with Σ1278b for 96 hours, larger, confluent areas of biofilm formed around the rough outer edge of the PET coupon (figure 4D), demonstrating that such biofilms can be stable and remain viable over several days.

**Figure 4.**
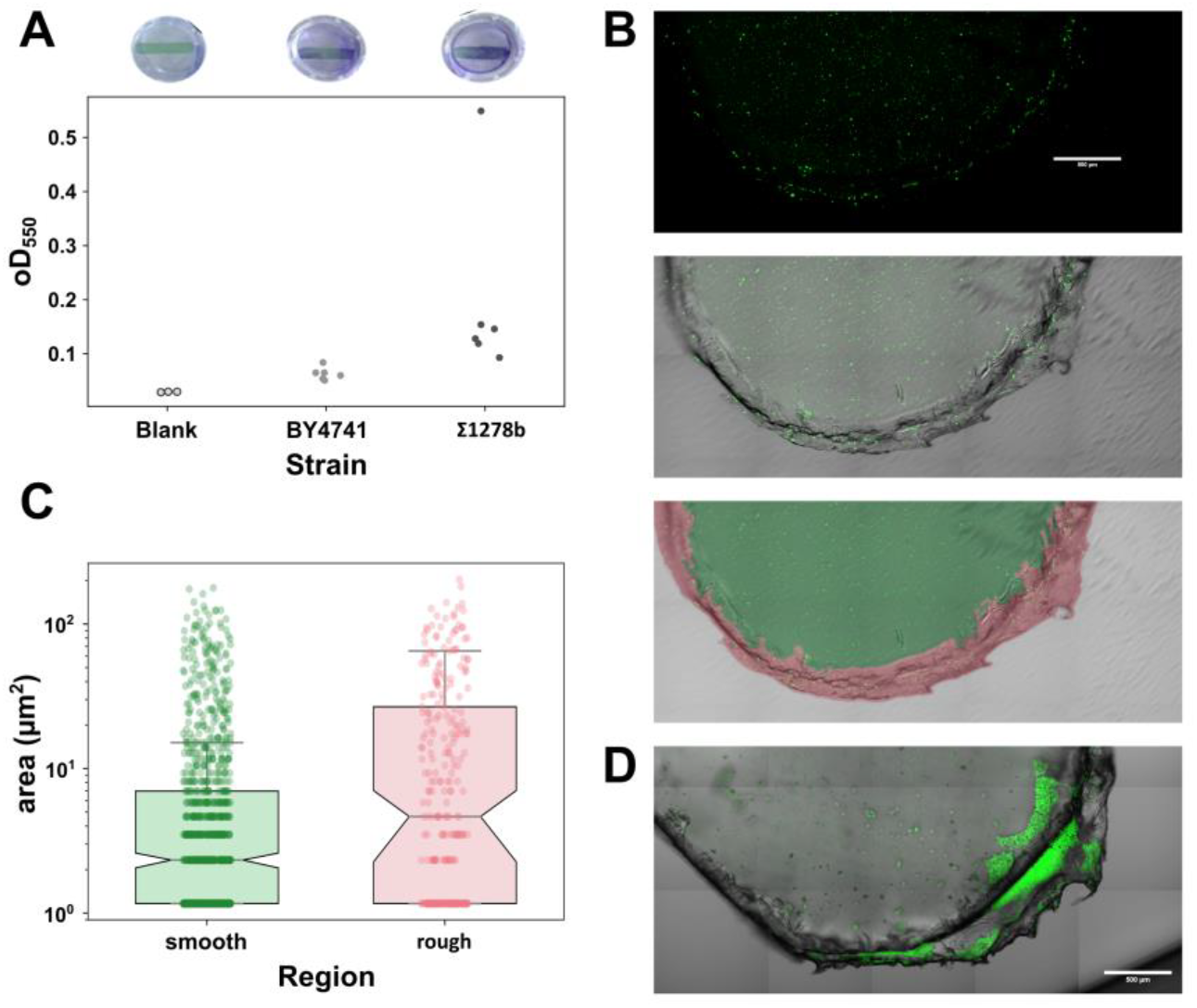
Σ1278b forms biofilms on PET substrates. **A**, Comparison of the ability of BY4741 and Σ1278b strains to adhere to PET coupons using a crystal violet assay. **B**, Biofilm formation on PET coupons following 24 hours of incubation. Cells were stained with Calcofluor White (CFW) and imaged. Top, confocal CFW image, middle, DIC image with overlaid image from the top panel, bottom, as middle image with indication of the “smooth” (green) and “rough” (red) zones used for colony size quantification. **C**, comparison of colony sizes in the red and green zones in panel B. **D**, CFW stained biofilm after 96 hours of incubation. Scale Bars, 500 μm.

A potential issue with the use of biofilms to degrade plastic is that yeast media are naturally slightly acidic, and yeast further acidifies growth media during the culture cycle (56). In contrast, commonly used PET hydrolases are most active under alkaline conditions, with optima around pH 8-9 (13). Yeast cells tolerate media that are buffered to neutral pH well, and indeed show longer chronological life span in such buffered media (57). To explore whether it was possible to buffer yeast cultures to a pH where *Is*PETase had higher activity, we tested yeast growth rates in media buffered to different target pH (supplemental figure 2). From pH 7 to pH 8, we observed that growth rates showed a moderate decline, but yeast grew actively and reached similar biomass as in unbuffered medium. At higher pH (>8.5), yeast appeared to show a very long lag phase, before eventually resuming growth. In the following we therefore conducted degradation assays in media buffered to pH8, which is both compatible with efficient yeast growth and high *Is*PETase activity.

### Degradation of PET by IsPETase-secreting yeast biofilms

To visualise PET degradation, we incubated coupons of amorphous PET film in growing cultures of *Is*PETase-secreting yeast strains. Following 96 hours of growth, PET surfaces were washed to remove attached yeast cells and visualised using two different imaging approaches, AFM (figure 5A and B) and SEM (figure 5C). Coupons incubated with yeast containing a control vector showed smooth surfaces with only few surface abrasions that appeared mechanical in origin (figure 4A). In contrast, coupons incubated with *Is*PETase-secreting yeast showed widespread changes to the PET surface indicative of PET degradation activity. These surface changes were less extensive on coupons incubated with the non-biofilm forming BY4741 strain, where they appeared as a general surface roughening in AFM images (figure 5A), and as small irregular holes in SEM images (figure 5C). Incubation with *Is*PETase secreting, biofilm forming Σ1278b resulted in more extensive surface changes visible as strong pitting in both imaging modes. We quantified these results by determining the surface roughness in our AFM images, confirming quantitatively that incubation with the biofilm-forming Σ1278b strain produced much stronger surface changes than the non-biofilm forming BY4741 (figure 5B and supplemental table 1).

**Figure 5.**
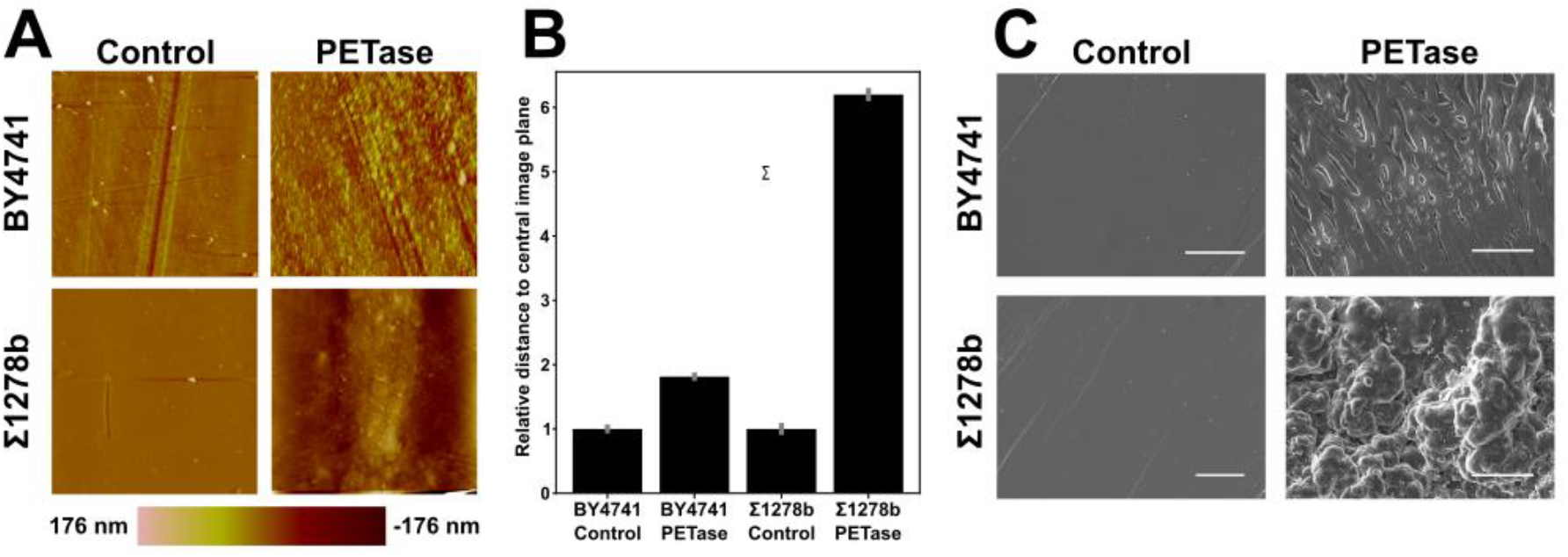
PET surface integrity following Incubation with PET secreting yeast strains. **A**, AFM images of control PET surfaces and surfaces following 24 hour incubation with PET hydrolase secreting yeast. Image areas are 20×20 μm. **B**, Quantification of average surface roughness in AFM images following biofilm growth of *Is*PETase secreting or control strains, based on the Ra5 measure (equation 1 in the Materials and methods section). **C**, SEM images of control PET surfaces and surfaces following 24 hour incubation with PET hydrolase secreting yeast. Scale bars are 15 μm.

As further evidence that yeast-secreted PET hydrolases are active, we sought to detect degradation products in the yeast growth medium. PET degradation results in a mixture of the monomeric subunits TPA and ethylene glycol (EG), as well as the condensation adducts mono and bis(2-hydroxyethyl)terephthalic acid (MHET and BHET, respectively). TPA and MHET are the predominant products detectable by HPLC, with BHET detectable at lower amounts (13). We were unable to procure MHET standards for HPLC assays, and therefore developed assays for TPA and BHET which are available commercially in purified form.

Pure TPA and BHET were suspended in spent yeast growth medium, and generated clear additional peaks in HPLC spectra of spent medium (supplemental figure 3), with the peak volume correlating with the amount of both chemicals spiked into the yeast media (figure 6). Peaks of similar retention times were also observed with media in which *Is*PETase secreting yeast had been grown in the presence of PET, but not when non-PET hydrolase secreting yeast were grown in the presence of PET (figure 6). Peaks in the PET degradation samples appeared to show random shifts in elution time, within a range of +/- 0.2 minutes around the standard samples (*c*.*f*. the lower panels in figure 6). These shifts may be related to pH shifts in the media containing growing yeast, since changes in an analyte’s ionisation state can strongly influence its exchange between the mobile and stationary phases (58). Even in our Tris-buffered media, the pH at the end of the incubation periods varied strongly (between 5 and 7), and differences in elution time in different samples may be related to these pH differences.

**Figure 6.**
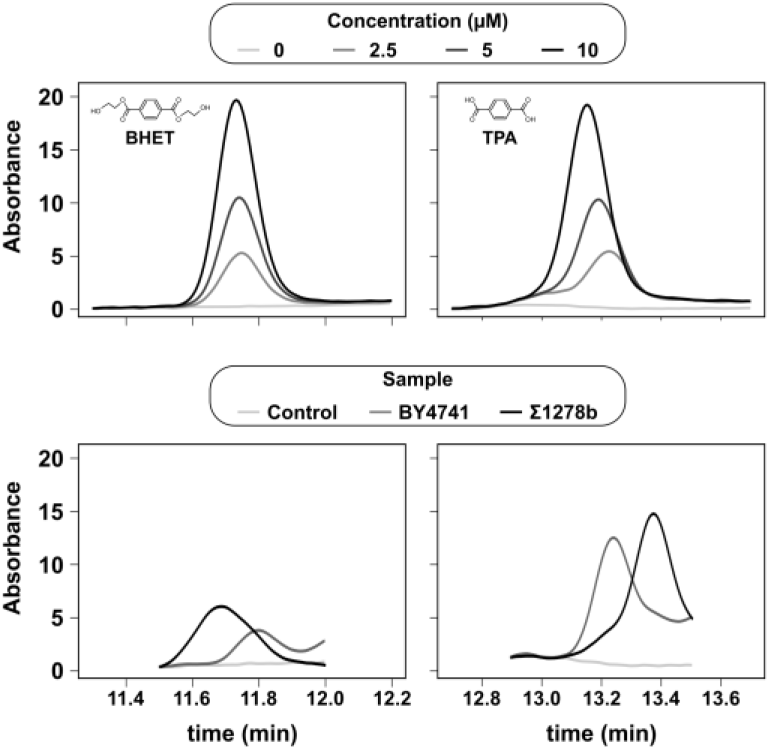
Detection of PET degradation products following incubation of PET coupons with PET hydrolase secreting yeast. Top, reference HPLC chromatograms of varying concentrations of the PET hydrolase products BHET and terephthalic acid (TPA). Bottom, HPLC analysis of media supernatants of non-PETase secreting yeast (light grey), PETase secreting BY4741 (dark grey) and PETase secreting Σ1278b (black).

In principle the apparent peak volumes for BHET and TPA can be used to estimate how much product was formed on our degradation assays. However, since BHET hydrolyses spontaneously to MHET and EG in aqueous solution, and since we were not able to develop an assay for MHET, quantitation based on our data is only partial. At the point of analysis following 96 hours of incubation, the yeast media contained 7.5 μM TPA and 3 μM BHET when incubated with Σ1278b, and 6 μM TPA and 1.8 μM BHET for BY4741. These analyses confirm the higher degradation activity for the biofilm forming Σ1278b strain, although the difference between the strains appears less than in the surface-based assay (figure 5B). Accounting for concentration steps during sample preparation and the amount of plastic suspended in the culture (each coupon contained about 100 mg PET), this corresponds to a conversion rate of 5 μM TPA and BHET per gram PET for the Σ1278b strain. In optimised systems and with purified enzymes, conversion rates of up to 5 mM total aromatic substances (TPA+MHET+BHET) per gram PET, albeit with very high suspended PET concentrations of 29g/l (59). We conclude that while our biofilm-based system in its current state is not quantitatively competitive with mature systems based on purified enzymes, it shows promising activity that could be further developed by optimising expression levels, biofilm forming activity, and specific enzyme activity.

In a final degradation assay, we compared the activity of different PETases against a model substrate (amorphous PET film) and against post-consumer PET samples (figure 7). In these AFM-based surface assays we could detect activity against both substrates, albeit with quantitatively differing preferences for different PET hydrolases. In summary, we provide evidence of activity of biofilm-delivered PET hydrolases against different PET substrates. Based on the current strains and enzymes evidence for degradation activity is at the microscopic level. These results nevertheless provide proof of principle that biofilm-based approaches can function in principle, and suggest possible routes towards optimisation of such approaches (see discussion for details).

**Figure 7.**
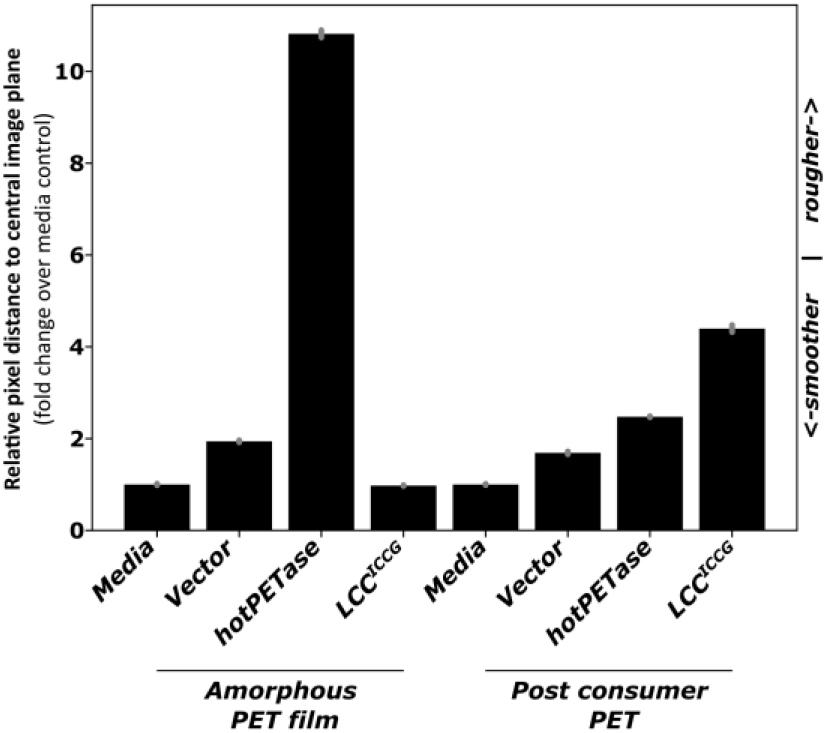
Comparison of the degradation activity of different biofilm-delivered PET hydrolases on a model substrate (amorphous PET film) and on post-consumer PET samples. PET hydrolase secreting strains or control strains were grown on PET samples for 96 hours. PET samples were washed to remove cells and imaged using AFM. Changes in surface roughness were then determined based on the average pixel distance from the central image plane in the AFM images.

### Yeasts can assimilate PET-derived degradation products

The products of the *Is*PETase reaction are small, carbon-containing entities which could potentially be assimilated by yeast. We assembled yeast media containing various PET degradation products as the sole carbon source, and tested whether yeast could form biomass in such media. These experiments are complicated by the fact that standard laboratory yeast strains require amino acid supplementation, and amino acids can be efficient carbon sources in their own right. We therefore conducted growth assays with a completely auxotrophic wild-type yeast strain, S288c (the parental wild-type strain of BY4741,(41)), which does not require any amino acid supplementation. In pure yeast nitrogen base only supplemented with DMSO (the vehicle used to dissolve stock solutions of BHET and TPA), yeast only showed a residual increase in cell numbers after 48 hours of incubation, presumably enabled by intracellular carbon stores (figure 8A and B). In contrast, in the presence of 0.5% BHET, TPA or EG, cell numbers following incubation were increased, and this increase was only statistically significant for the BHET sample (figure 6A). Because different carbon sources can affect yeast cell size and this can confound optical density measurements (60), these assays were conducted by counting cell numbers using hemocytometers. In addition, we were also able to observe growth in the presence of BHET as sole carbon source when we monitored culture oD continuously over time (figure 8B).

**Figure 8.**
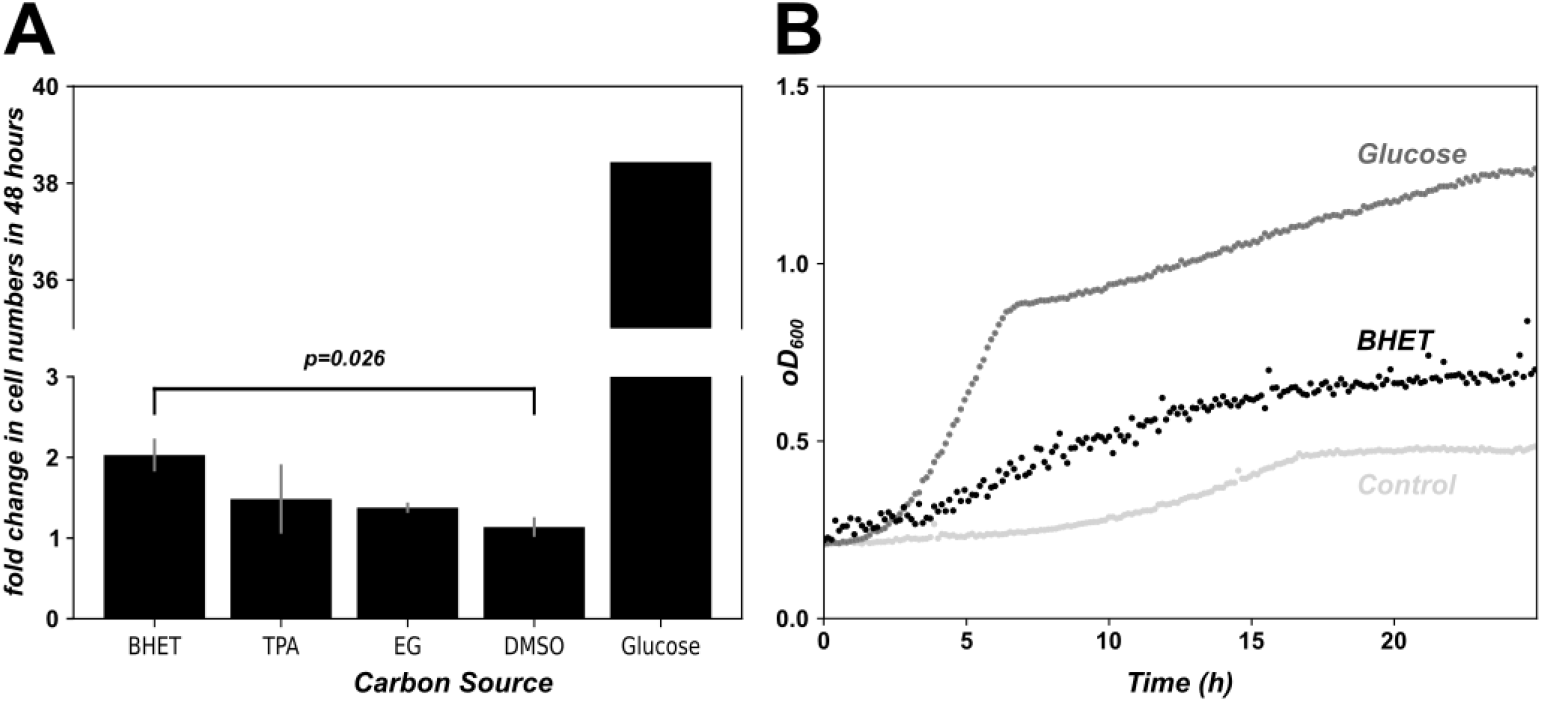
Yeast can assimilate PET degradation products into biomass. **A**, *S. cerevisiae* S288c (the wild-type parental strain of BY4741) were inoculated into synthetic defined medium containing either glucose or various PET degradation products as sole carbon source. All samples were adjusted to contain equal amounts of DMSO (the solvent used to prepare BHET and TPA stock solutions). The increase in microscopically determined cell numbers following 48 hours of incubation is shown. Feathers indicate the standard error of the mean (n=5). **B**, growth curves for the same yeast strain grown in the presence of glucose, BHET or no added carbon source (samples were adjusted to equal DMSO concentration as in A).

In conclusion, these results indicate that yeast can take up and assimilate either BHET or a spontaneous hydrolysis product of BHET (our assays do not allow us to distinguish between these possibilities). In either case, we demonstrate that treatment of PET with PET hydrolase enzymes generates bioavailable products, and that *Is*PETase producing biofilms are thus a potentially useful tool for the bioremediation of PET-contaminated soils and other environments.

## Discussion

PET hydrolases have been expressed in a number of heterologous expression systems (table 1) which are optimised for different purposes. The production of bulk enzyme for purification is a frequent purpose, typically with the aim of fully recycling PET waste. For this application high expression levels are typically required to achieve commercial viability. Other, more specialised applications that have been proposed include the development of degrading consortia that facilitate conversion of the degradation products into biomass (47)(44), and the conversion of the PET degradation products into high value compounds such as vanillin (62).

Enzyme-based applications proposed or developed to date are predominantly tailored to the treatment of clean plastic. Plastic recovered from environmental sources frequently contains mixed plastic chemistries as well as other impurities, and the chemical structure of such recovered plastic is often altered due to the influence of UV light, oxygen, and other factors [ref] (63). The ability to recycle such plastic waste through melting and reshaping is limited, as the chemical changes in the plastic lead to a loss of tensile strength (64). We are not aware of studies that have directly addressed how recyclable such waste is with enzyme-based approaches, but the presence of impurities would presumably affect the ability to recover clean monomeric subunits that are amenable to the re-polymerisation of virgin plastic. Operations to recover environmental plastic are a growing activity area with operations such as the Ocean Cleanup and Shoreline Cleanups (65) now recovering tens to hundreds of tons of plastic waste annually. Biofilm-based solutions could play a significant role in the removal of such recovered waste.

Our current study demonstrates the delivery of degrading enzymes close to a target surface can enhance the degree of degradation. Based on the increase in surface roughness, we estimate that PET is removed at least 3 times more efficiently by a biofilm-forming PET hydrolase-secreting strain compared to a planktonic one (figure 4B). Our *S. cerevisiae*-based approach so far only delivers microscopic evidence of degradation and does not yet allow full deconstruction of plastic waste, but our work defines specific parameters that need to be improved for a functioning bioremediation solution.

With the strains selected for our study we observed limited secretion levels. This is common for unoptimised baker’s yeast strains, but expression levels can frequently be improved through random or rational strain engineering approaches (66). One successful approach for disulfide-bonded proteins is the overexpression of elements of the oxidative folding machinery (67), although in our hands overexpression of neither yeast *PDI1* nor of *ERO1* significantly improved *Is*PETase secreation (supplemental figure 4). Other optimisation approaches include modification of gene copy numbers or co-expression of chaperones, although we did not explore secretion optimisation further in this study.

Work published with purified enzymes indicates that degradation is optimal at ratios above 2 mg enzyme per gram plastic at PET loads of 29g per litre. For planktonic cells where the enzyme is roughly evenly distributed in the growth medium, secretion would need to be improved by several orders of magnitude, to levels of 50-100 mg/l, to achieve such ratios. Inside biofilm-matrices macromolecules remain trapped with half-lives of minutes (36), and this would thus significantly increase the local enzyme concentration at the target surface over the average concentration in the overall culture, reducing the requirement for increased expression levels.

Our observation that *Is*PETase is glycosylated, but that the degree of glycosylation is reduced with engineered forms of the protein that fold more stably (figure 3), indicates that the ability to fold correctly may be limited at least for *Is*PETase (LCC did not appear to be significantly glycosylated in baker’s yeast). *Is*PETase secreted from *Pichia pastoris* cells was similarly observed to be glycosylated, and partial enzymatic removal of the glycan chains was shown to increase enzyme activity (68). Thus glycosylation is likely another parameter that needs to be controlled for efficient degradation. One approach to achieve this could be the removal of surface-accessible asparagine residues in the enzymes, although if the glycosylation is a response to incorrect folding of the enzyme as our observation that more stably folding mutants are less glycosylated suggests, this would adversely affect expression levels and host cell health. Since Leaf Compost Cutinase appears to be expressed without significant glycosylation (figure 3), as well as being the most active enzyme against post-consumer waste (figure 7 and supplemental table 1), this enzyme would be a natural starting point for further development of biofilm-based degradation systems.

Overall, our results show that engineered biofilms are a promising approach for delivering plastic-degrading enzymes onto plastic surfaces and, with further development, could yield useful tools for the bioremediation of environmental plastic.

## Methods

### Plasmid Construction and Codon optimisation

The native PET hydrolase genes contain codons which are rarely used and suboptimal in baker’s yeast (Figure 1A). To increase the probability of observing high expression levels in yeast and ensure secretion of the expressed protein into the extracellular medium, we designed yeast optimised DNA sequences encoding N-terminal secretion signal sequences from the yeast *SUC2* gene fused to different PET hydrolase genes, followed by C-terminal HA tags to facilitate detection (Figure 1A). Synthetic genes encoding these proteins were introduced into pTH644 (38), a single-copy yeast expression vector, under control of the strong constitutively active promoter of the yeast *TDH3* gene.

The combined amino acid sequences were then converted into DNA sequences by selecting the fastest decoded codons in *S. cerevisiae* for each amino acid as described (39). Optimised gene sequences are listed in the supplemental data, and physical sequences were synthesised by GenScript (UK). The PETase genes were amplified by polymerase chain reaction (PCR, primer sequences are listed in supplemental table 2), introducing overlaps for insertion into *HindIII* / *BamHI* linearised pTH644 (38) using Gibson assembly (40). All PET hydrolase expression plasmids are available from Addgene ^1^ (supplemental table 3).

### Yeast strains and growth conditions

*S. cerevisiae* strains used in this study include BY4741 (MAT**a** *his3Δ1 leu2Δ0 met15Δ0 ura3Δ0*) (41), Σ1278b (MATα *ura3-52; trp1Δ::hisG; leu2Δ::hisG; his3Δ::hisG*) (42) and S288c (MAT **a**) (43).

Yeast cells were transformed using the lithium acetate method (44). Transformants were grown in selective conditions using Synthetic Dropout media lacking uracil (SC-Ura, 20 g/l glucose, 6.7 g/l Yeast Nitrogen Base without amino acids, 1.9 g/l Amino Acid Dropout Supplement without uracil (Formedium, UK)). Where indicated the pH of the media was adjusted by adding Tris-HCl buffer to a final concentration of 20 mM from 1M stock.

For analyses of growth rates transformed cells were grown in selective medium in 24 well Greiner plates. The plates were incubated in a BMG Labtech (UK) SPECTROstar microplate reader, and absorbance at 600nm was recorded every 8 minutes.

### Protein detection and purification

For analyses of PET hydrolase expression, cells were grown overnight in selective medium and 1ml of the culture was centrifuged at 8,000 rpm for 5 minutes. Supernatant and cell pellet were recovered and analysed separately.

For intracellular expression, cell extracts were prepared as described (45). For analyses of secreted protein, the supernatant was concentrated by adding 100 μl 0.15% (w/v) sodium deoxycholate and 50μL of 100% trichloracetic acid (TCA), incubation on ice for 10 minutes, and centrifugation in a microcentrifuge at 13,000 rpm for 20 minutes. Pellets were washed twice with ice-cold acetone, dried and resuspended in 50 μL of 2x Laemmli Sample Buffer, containing 10% (v/v) β-mercaptoethanol.

Both the cell lysate and the supernatant samples were separated by sodium dodecyl sulfate polyacrylamide gel electrophoresis (SDS-PAGE) and Western blotting as described (45) using antibodies from Sigma-Aldrich, UK (anti-HA tag, H6908, peroxidase-labelled anti-rabbit IgG, A9169). For analyses of glycosylation, growth supernatant was treated with Endo Hf (NEB, UK) as directed by the manufacturer prior to blotting.

For the quantification of secreted HA-tagged proteins, we added a C-terminal HA tag to the glutathione-S-transferase (GST) encoded on the pGEX-6P-1 vector (GE Healthcare, UK). HA-tagged GST was expressed in *E. coli* BL21 cells and purified using chelating sepharose 4B (Sigma Aldrich, UK) according to the manufacturer’s instructions. HA-tagged GST protein was used a a reference for determining amounts of other HA-ytagge dproteins on gels using quantitative western blotting as described (46).

### PET degradation assays

PET coupons were made from either amorphous PET film (2-3% crystallinity, Goodfellow, UK) or from post-consumer plastic items which were clearly labelled with the PET symbol using a 5 × 3 mm Metal Punch. Transformed *S. cerevisiae* cells were grown in 50 mL glass tubes with 5ml of SC-Ura pH 8.0 at 30°C with 180 rpm agitation for up to 1 week. Each glass tube contained 1 PET coupon. All cell cultures were inoculated to a starting cell density of 1.5x 10^8^ cells/ml. Before inoculation and after incubation PET coupons were cleaned using consecutive washes with 10% (w/v) SDS, 70% (v/v) ethanol, and dimethyl sulfoxide (DMSO), rinsing with deionised water in between washes. Following a final rinse with deionised water samples were left to dry at 70°C for 30 minutes.

Washed and dried coupons were then analysed for degradation by Scanning Electron Microscopy (SEM) and Atomic Force Microscopy (AFM). For SEM, the PET coupons were mounted onto 25 mm aluminium SEM specimen stubs (Agar Scientific, AGG3377) with 12 mm carbon adhesive tabs (Agar Scientific, AGG3347N). Samples were sputter coated with carbon using a Q150T Plus Turbomolecular pumped coater (Quorum, UK). The surfaces were imaged at room temperature using a Hitachi S-3400N in SEM mode with a chamber pressure of 0 Pa and an accelerating voltage of 5 kV. The backscattered electron detector was used with a working distance of 10 nm.

For AFM, the PET coupons were attached to 15 mm specimen discs with 12 mm carbon adhesive tabs (both from Agar Scientific, UK). Topology imaging was undertaken in air on a Bruker multimode 8 scanning probe microscope with a Nanoscope V controller, using the ScanAsyst peak-force tapping image mode. RTESPA-300 cantilever probes with a nominal spring constant of 40 N/m and a nominal tip radius of 8 nm (Bruker, Germany) were used for all analyses. Height images were captured at 20 × 20 μm scan size at 1024×1024 pixels or 40 × 40 μm scan size at 2048×2048 pixels. Images were then processed to remove scanner tilt and bow using Nanoscope analysis 1.5 (Bruker, Germany). The roughness of the imaged sample surfaces was quantified and compared using the height topology image channel by calculating the average deviation, R_a_ (Eq. 1) and root mean square deviation, R_q_ (Eq. 2) of surface heights from a mean plane. For this analysis, a series of 5 um image crops were each fit to a first order polynomial plane before the surface roughness R_a_ and R_q_ parameters and their standard errors were calculated.

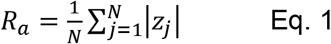

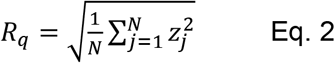

In Eq. 1 and 2, z are the pixel height values relative to the mean plane and N is the number of pixels.

### Detection of PET degradation products by high-performance liquid chromatography (HPLC)

PET samples were incubated with growing yeast cells as stated above for SEM and AFM analyses. Growth cultures were then prepared for HPLC analysis by initially removing the plastic coupon from the transformed cells, followed by centrifugation at 4,000 rpm for 20 minutes. 1 ml of the supernatant was recovered, placed into 2 ml microcentrifuge tubes, and frozen on dry ice. After incubation overnight at -80°C samples were freeze dried using an Edwards Modulyo freeze dryer for 16 hours. Prior to analysis samples were resuspended in 50 μl sterile H_2_O. Standards at the following concentrations: 500, 100, 50, 25, 10, 5 and 2.5 μM, were prepared for both terephthalic acid (TPA) and bis(2-hydroxyethyl) terephthalate (BHET, both from Sigma Aldrich, UK).

Samples were analysed by HPLC. Analytes were detected using a diode array detector collecting signals at 240 nm following separation on an XTerra RP18 column (3.5 μM resin bead diameter, 4.6 × 150mm, Waters, UK) using 20mM phosphoric acid in water (Solution A) and 100% methanol (Solution B). Separation was carried out at 30°C with a flow rate of 0.6 ml/min and the following gradient of Solution A/Solution B: 0-15 minutes 80%/20%, 15-20 minutes 35%/65%, 20-25 minutes 0%/100%.

### Biofilm formation analysis

For initial analyses, post-consumer PET coupons were placed in yeast cultures of 1 ml SC-Ura buffered with 20 mM Tris-HCl to pH 8.0 inoculated with *S. cerevisiae* cells to a density of 1.5 × 10^8^ cells per ml in a 24 well plate. Each strain of *S. cerevisiae* was grown with a PET coupon for 24-48 hours at 30°C shaking at 180 rotations per minute (rpm). After planktonic cells were removed by aspiration, plastic samples with remaining attached cells were incubated with 0.1% (w/v) crystal violet solution for 15 minutes and were then subjected to 2 washes in sterile water before being dried at 70°C for 1 hour. Images were taken once more of the bottom of the wells using an Epson scanner. Finally, images of the plastic with cells stained with crystal violet were acquired using Leica MZFLIII Fluorescence Stereozoom Microscope with image acquisition using a CellCam Rana 200CR colour camera. The images were processed using ImageJ and exported as a JPEG file. To quantify washing-resistant, crystal violet stained cells, PET coupons were then transferred into a new 24 well plate containing 30% acetic acid and incubated at room temperature for 15 minutes. Following this 100 μL of the solubilised crystal violet was transferred to a new 96 well plate where absorbance was measured at 550 nm on the BMG Labtech Spectrostar Omega Microplate Reader.

For higher magnification imaging, PET coupons after 24 hours or 48 hours of incubation with growing yeast cells in SC-Ura medium were imaged using a Zeiss LSM880 Elyra Confocal Microscope (Carl Zeiss Inc.). Biofilms on PET coupons were washed once with distilled water and stained with calcofluor white for 5 minutes before imaging. Images were taken with a 10x Plan-Apochromat (NA 0.45) M27 objective lens. Yeast cells were visualised using Calcofluor White, excited at 405 nm and collected at 450/40 emission. Differential interference contrast (DIC) images were collected using 488 nm excitation and a transmitted light PMT detector. Tile-stitched Z stacks were captured to visualise the full end of a coupon in all three spatial dimensions. Zeiss ZEN software (black edition) was used for image acquisition. ImageJ 1.53t was used to quantify images, with Otsu thresholding in the Calcofluor White channel to identify yeast colonies, followed by “Analyze Particles” to determine the colony sizes. The coupon was divided into rough and smooth zones based on visual inspection of the DIC image.

## Data analysis and data availability

Data were analysed using Python 3.7.11, pandas 1.3.4, numpy 1.21.6, matplotlib 3.4.3 and statsmodels 0.12.2.

## Data availability

All data and analysis scripts are available for download^2^.

## Acknowledgements

This work was supported by a PhD studentship funded by the Biotechnology and Biological Sciences Research Council (UK) through the South Coast Biosciences Doctoral Training Partnership (to CRB), BBSRC grant BB/W011530/1 (to WFX) and by a Leverhulme Research Fellowship (RF-2019-455, to TVDH).

## Author contributions

CRB, JD, WFX, CG, PPL, ARP, AL and TVDH designed the study. CRB, NI and TVDH generated reagents. CRB, SKH, JD, SMJ, PPL, EDR and SS conducted experimental analyses. CRB, SKH, JD, SMJ, RG, EDR, SS, WFX, AL and TVDH analysed data. WFX, ARP and TVDH secured funding. All authors read and approved the manuscript.

## Footnotes

1 https://www.addgene.org/plasmids/articles/28238320/

1 github.com/tobiasvonderhaar/PETaseBiofilms

