## Supplemental materials for "Engineered yeast biofilms deliver plastic degrading enzymes to PET substrates"

### Supplemental Materials for Bilsby *et al.* (2025)

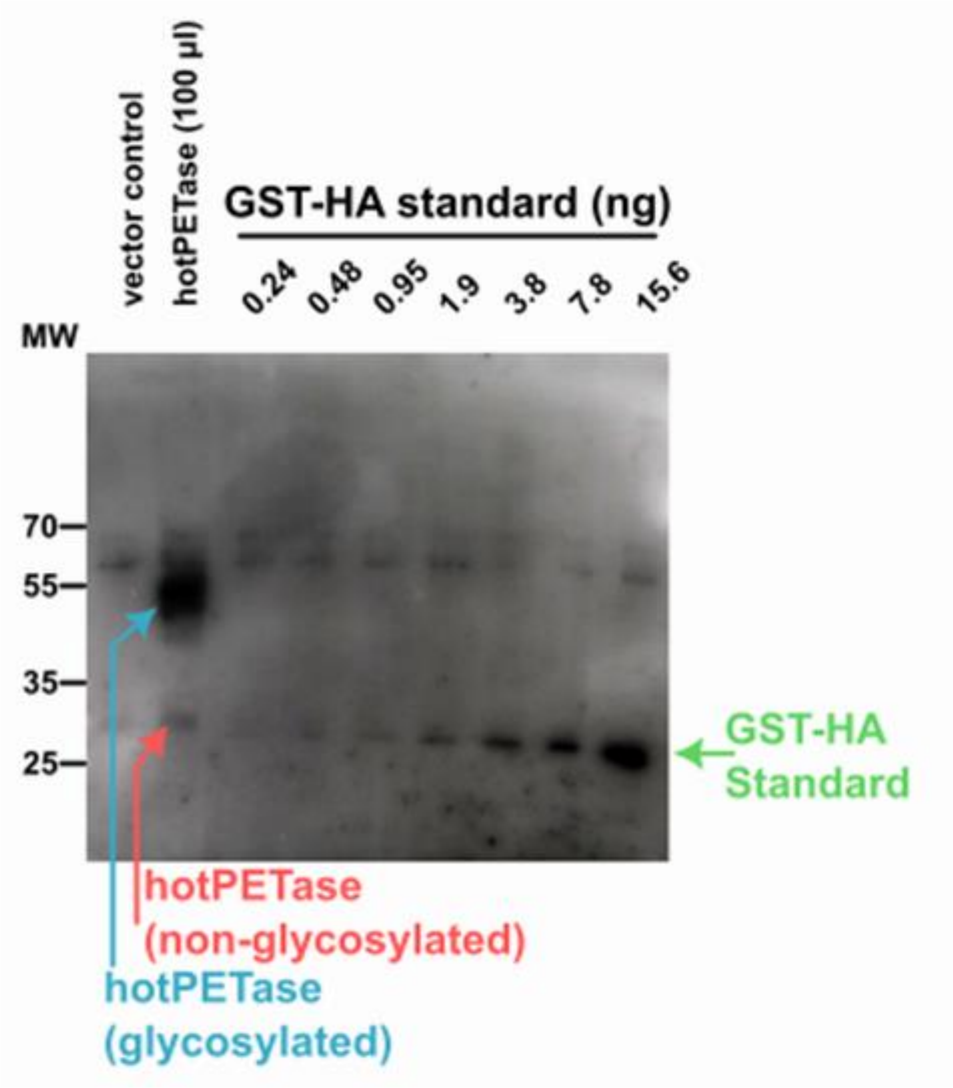

**Supplemental figure 1. Quantitation of HA-tagged hotPETase against quantitative standards of HA-tagged GST protein.** See main text for explanation.

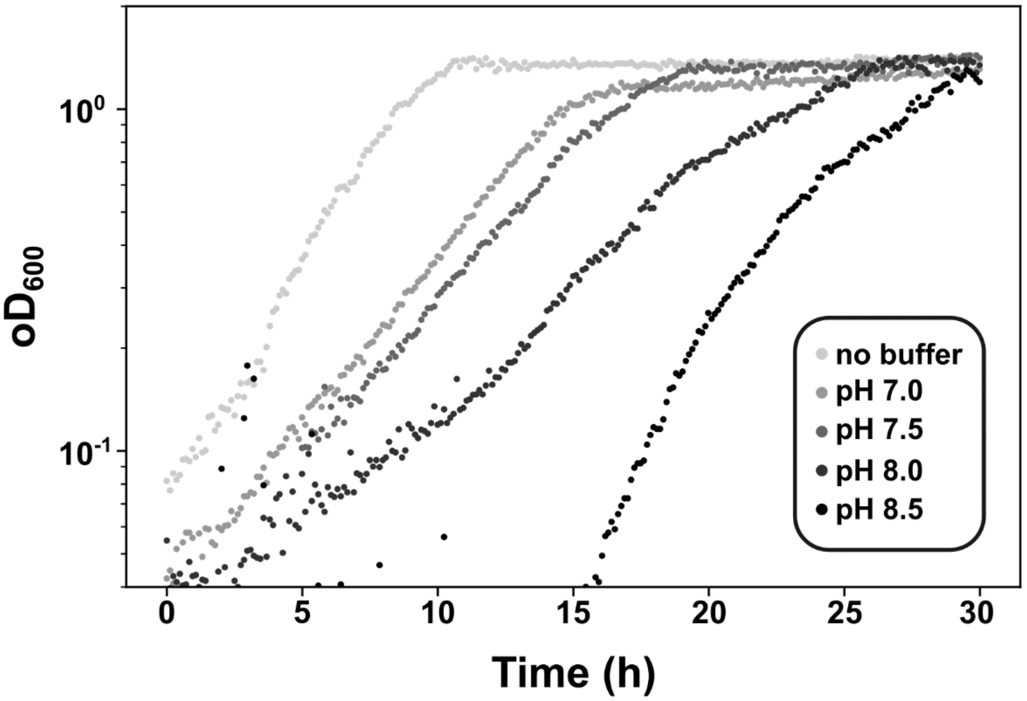

**Supplemental figure 2. Yeast growth in pH buffered media.** The ability of yeast to grow in pH buffered media that match the alkaline pH optimum of PET hydrolase was assessed.

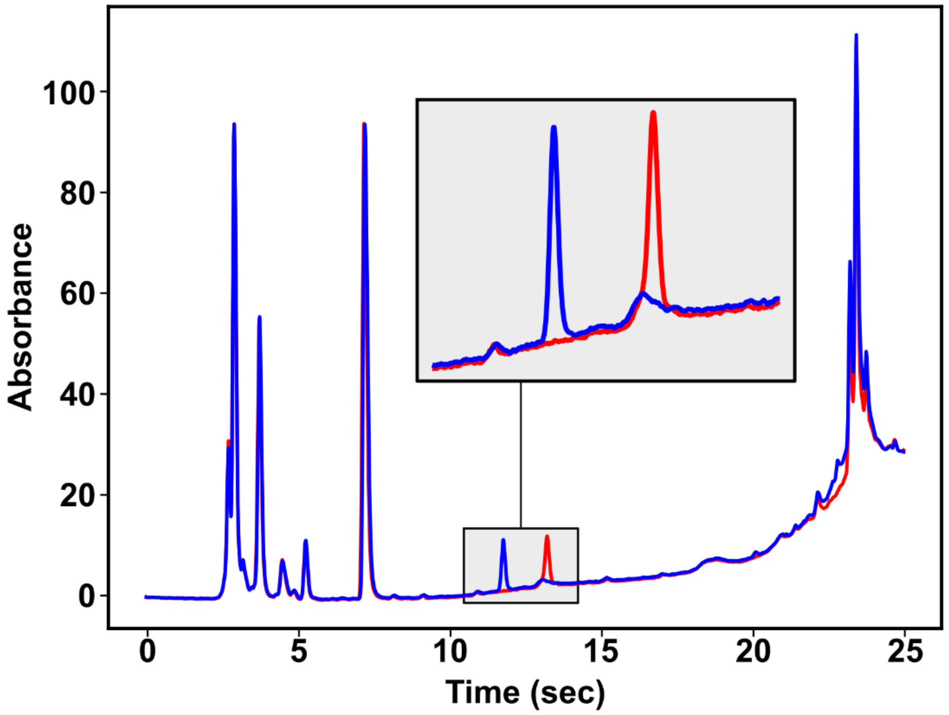

**Supplemental Figure 3.** Full HPLC chromatograms for BHET (Blue) and TPA (red) standards (5 µM in spent SC -Ura medium).

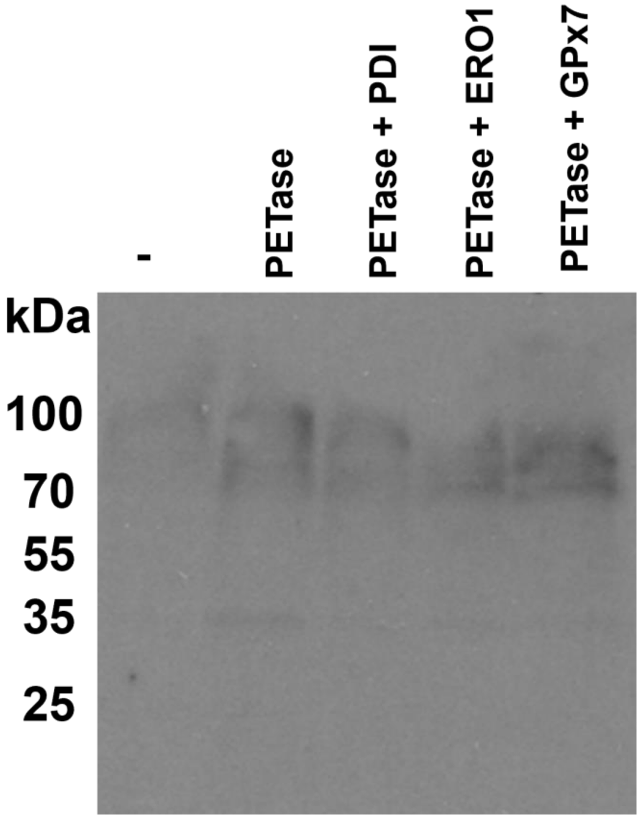

**Supplemental figure 4. *Is*PETase expression levels in the presence of co-expressed oxidative folding factors.** Overexpression of the native yeast protein disulfide isomerase, Pdi1p, of the native yeast ER Oxidoreductin, Ero1p, or of human GPx7 can affect the expression of some proteins containing disulfide bonds, but did not affect expression levels of *Is*PETase. The blot was processed using anti-HA primary antibodies which detect the C-terminal HA tag present in our *Is*PETase expression constructs.

**Supplemental table 1.** Surface roughness data.

| | Figure 5 | | | | | | |
|---|---|---|---|---|---|---|---|
| | | | Ra_5 | SE | | Rq_5 | SE |
| | BY4741 control | | 1.00 | 0.16 | | 1.00 | 0.27 |
| | BY4741 PETase | | 2.34 | 0.31 | | 1.85 | 0.41 |
| | ∑1278b control | | 1.00 | 0.06 | | 1.00 | 0.31 |
| | ∑1278b PETase | | 5.08 | 0.23 | | 2.79 | 0.55 |
| Amorphous PET | media control | | 1.00 | 0.08 | | 1.00 | 0.18 |
| | vector | | 1.95 | 0.17 | | 1.86 | 0.32 |
| | hotPETase | | 8.88 | 1.40 | | 6.89 | 1.95 |
| | LCC_ICCG | | 1.18 | 0.07 | | 0.97 | 0.12 |
| Post Consumer PET | media control | | 1.00 | 0.07 | | 1.00 | 0.15 |
| | vector | | 1.69 | 0.13 | | 1.63 | 0.33 |
| | hotPETase | | 2.37 | 0.17 | | 2.03 | 0.27 |
| | LCC_ICCG | | 3.39 | 0.38 | | 3.93 | 0.93 |

**Supplemental table 2.** Primer sequences. All primers were synthesized by Integrated DNA technologies (USA).

| Name | Sequence | Purpose |
|---|---|---|
| PET-f | TCGAGGTCGACGGTATCGATAAGCTTGCTTTTCATAGGGTAGGGGAATTTC | *Is*PETase cloning into pTH644 |
| PET-r | GGGCTGCAGGAATTCGATATC*AAGCTT*CATTATCAATACTGCCATTTCAAAGAA | |
| opt_gibson_f | GTTTTAAAACACCAAGAACTTAGTTTCGACGG | Cloning of synthetic genes synthesized with compatible overhangs into pTH644 |
| opt_gibson_r | CTTGACCAAACCTCTGGCGAAGAAGTCC | |

**Supplemental table 3.** Plasmids used in this study.

| Plasmid | Insert | Purpose | Reference | Addgene No |
|---|---|---|---|---|
| pTH644 | - | Single copy bidirectional expression vector for baker's yeast | Chu *et al.*, 2011 | 29695 |
| pCB11 | Yeast codon optimised *Is*PETase fused to an Ost/MAT signal sequence and C-terminal HA tag | Expression and secretion of *Is*PETase | This study | 205605 |

| | | | | |
|---|---|---|---|---|
| pCB13 | Yeast codon optimised *Is*PETase$^{Ts}$ fused to a *SUC2* signal sequence and C-terminal HA tag | Expression and secretion of *Is*PETase$^{Ts}$ | This study | 211808 |
| pCB36 | Yeast codon optimised hotPETase fused to a SUC2 signal sequence and C-terminal HA tag | Expression and secretion of hotPETase | This study | tbc |
| pCB16 | Yeast codon optimised Leaf Compost Cutinase fused to a *SUC2* signal sequence and C-terminal HA tag | Expression and secretion of LCC | This study | 211809 |
| pCB35 | Yeast codon optimised, engineered Leaf Compost Cutinase (LCC$^{ICCG}$) fused to a *SUC2* signal sequence and C-terminal HA tag | Expression and secretion of LCC$^{ICCG}$ | This study | tbc |

**Supplemental Material - Optimised Gene Sequence**

```
>scopt_IsPETase

ATGTTGTTGCAAGCGTTTTTGTTTTTGTTGGCGGGTTTTGCGGCGAAAATTTCCGCGTCCATGCAAAC
TAACCCGTACGCGAGAGGTCCGAACCCGACTGCGGCGTCCTTGGAAGCGTCCGCGGGTCCGTTTACTG
TCAGATCCTTTACTGTCTCCAGACCGTCCGGTTACGGTGCGGGTACTGTCTACTACCCGACTAACGCG
GGTGGTACTGTCGGTGCGATTGCGATTGTCCCGGGTTACACTGCGAGACAATCCTCCATTAAATGGTG
GGGTCCGAGATTGGCGTCCCATGGTTTTGTCGTCATTACTATTGACACTAACTCCACTTTGGACCAAC
CGTCCTCCAGATCCTCCCAACAAATGGCGGCGTTGAGACAAGTCGCGTCCTTGAACGGTACTTCCTCC
TCCCCGATTTACGGTAAAGTCGACACTGCGAGAATGGGTGTCATGGGTTGGTCCATGGGTGGTGGTGG
TTCCTTGATTTCCGCGGCGAACAACCCGTCCTTGAAAGCGGCGGCGCCGCAAGCGCCGTGGGACTCCT
CCACTAACTTTTCCTCCGTCACTGTCCCGACTTTGATTTTTGCGTGCGAAAACGACTCCATTGCGCCG
GTCAACTCCTCCGCGTTGCCGATTTACGACTCCATGTCCAGAAACGCGAAACAATTTTTGGAAATTAA
CGGTGGTTCCCATTCCTGCGCGAACTCCGGTAACTCCAACCAAGCGTTGATTGGTAAAAAAGGTGTCG
CGTGGATGAAAAGATTTATGGACAACGACACTAGATACTCCACTTTTGCGTGCGAAAACCCGAACTCC
ACTAGAGTCTCCGACTTTAGAACTGCGAACTGCTCCTACCCGTACGACGTCCCGGACTACGCGTAA

>scopt_IsPETase_Ts

ATGTTGTTGCAAGCGTTTTTGTTTTTGTTGGCGGGTTTTGCGGCGAAAATTTCCGCGTCCATGCAAAC
TAACCCGTACGCGAGAGGTCCGAACCCGACTGCGGCGTCCTTGGAAGCGTCCGCGGGTCCGTTTACTG
TCAGATCCTTTACTGTCTCCAGACCGTCCGGTTACGGTGCGGGTACTGTCTACTACCCGACTAACGCG
GGTGGTACTGTCGGTGCGATTGCGATTGTCCCGGGTTACACTGCGAGACAATCCTCCATTAAATGGTG
GGGTCCGAGATTGGCGTCCCATGGTTTTGTCGTCATTACTATTGACACTAACTCCACTTTGGACCAAC
CGGAATCCAGATCCTCCCAACAAATGGCGGCGTTGAGACAAGTCGCGTCCTTGAACGGTACTTCCTCC
TCCCCGATTTACGGTAAAGTCGACACTGCGAGAATGGGTGTCATGGGTTGGTCCATGGGTGGTGGTGG
TTCCTTGATTTCCGCGGCGAACAACCCGTCCTTGAAAGCGGCGGCGCCGCAAGCGCCGTGGCATTCCT
CCACTAACTTTTCCTCCGTCACTGTCCCGACTTTGATTTTTGCGTGCGAAAACGACTCCATTGCGCCG
GTCAACTCCTCCGCGTTGCCGATTTACGACTCCATGTCCAGAAACGCGAAACAATTTTTGGAAATTAA
CGGTGGTTCCCATTCCTGCGCGAACTCCGGTAACTCCAACCAAGCGTTGATTGGTAAAAAAGGTGTCG
CGTGGATGAAAAGATTTATGGACAACGACACTAGATACTCCACTTTTGCGTGCGAAAACCCGAACTCC
ACTGCGGTCTCCGACTTTAGAACTGCGAACTGCTCCTACCCGTACGACGTCCCGGACTACGCGTAA

>scopt_hotPETase

ATGTTGTTGCAAGCGTTTTTGTTTTTGTTGGCGGGTTTTGCGGCGAAAATTTCCGCGTCCATGCAAAC
TAACCCGTACGCGAGAGGTCCGAACCCGACTGCGGCGTCCTTGGAAGCGTCCGCGGGTCCGTTTACTG
TCAGATCCTTTACTGTCGCGAGACCGGTCGGTTACGGTGCGGGTACTGTCTACTACCCGACTAACGCG
GGTGGTACTGTCGGTGCGATTGCGATTGTCCCGGGTTACACTGCGACTCAATCCTCCATTAACTGGTG
GGGTCCGAGATTGGCGTCCCATGGTTTTGTCGTCATTACTATTGACACTAACTCCACTTTGGACAAAC
CGGAATCCAGATCCTCCCAACAAATGGCGGCGTTGAGACAAGTCGCGTCCTTGAACGGTACTTCCTCC
TCCCCGATTTACGGTAAAGTCGACACTGCGAGAGGTGGTGTCATGGGTTGGTCCATGGGTGGTGGTGG
TTCCTTGATTTCCGCGGCGAACAACCCGTCCTTGAAAGCGGCGGCGGTCATGGCGCCGTGGCATTCCT
CCACTAACTTTTCCTCCGTCACTGTCCCGACTTTGATTTTTGCGTGCGAAAACGACAGAATTGCGCCG
GTCAAAGAATACGCGTTGCCGATTTACGACTCCATGTCCTTGAACGCGAAACAATTTTTGGAAATTTG
CGGTGGTTCCCATTCCTGCGCGTGCTCCGGTAACTCCAACCAAGCGTTGATTGGTATGAAAGGTGTCG
CGTGGATGAAAAGATTTATGGACAACGACACTAGATACTCCCAATTTGCGTGCGAAAACCCGAACTCC
ACTGCGGTCTGCGACTTTAGAACTGCGAACTGCTCCTACCCGTACGACGTCCCGGACTACGCGTAA

>scopt_LCC
```

ATGGACGGCGTCTTGTGGAGAGTCAGAACTGCGGCGTTGATGGCGGCGTTGTTGGCGTTGGCGGCGTG
GGCGTTGGTCTGGGCGTCCCCGTCCGTCGAAGCGCAATCCAACCCGTACCAAAGAGGCCCGAACCCGA
CTAGATCCGCGTTGACTGCGGACGGCCCGTTTTCCGTCGCGACTTACACTGTCTCCAGATTGTCCGTC
TCCGGCTTTGGTGGTGGCGTCATTTACTACCCGACTGGCACTTCCTTGACTTTTGGTGGCATTGCGAT
GTCCCCGGGTTACACTGCGGACGCGTCCTCCTTGGCGTGGTTGGGCAGAAGATTGGCGTCCCATGGCT
TTGTCGTCTTGGTCATTAACACTAACTCCAGATTTGACTACCCGGACTCCAGAGCGTCCCAATTGTCC
GCGGCGTTGAACTACTTGAGAACTTCCTCCCCGTCCGCGGTCAGAGCGAGATTGGACGCGAACAGATT
GGCGGTCGCGGGCCATTCCATGGGCGGTGGTGGCACTTTGAGAATTGCGGAACAAAACCCGTCCTTGA
AAGCGGCGGTCCCGTTGACTCCGTGGCATACTGACAAAACTTTTAACACTTCCGTCCCGGTCTTGATT
GTCGGCGCGGAAGCGGACACTGTCGCGCCGGTCTCCCAACATGCGATTCCGTTTTACCAAAACTTGCC
GTCCACTACTCCGAAAGTCTACGTCGAATTGGACAACGCGTCCCATTTTGCGCCGAACTCCAACAACG
CGGCGATTTCCGTCTACACTATTTCCTGGATGAAATTGTGGGTCGACAACGACACTAGATACAGACAA
TTTTTGTGCAACGTCAACGACCCGGCGTTGTCCGACTTTAGAACTAACAACAGACATTGCCAATACCC
ATACGATGTTCCAGATTACGCTTAA

>scopt_LCC_ICCG

ATGTTGTTGCAAGCGTTTTTGTTTTTGTTGGCGGGTTTTGCGGCGAAAATTTCCGCGATGGACGGTGT
CTTGTGGAGAGTCAGAACTGCGGCGTTGATGGCGGCGTTGTTGGCGTTGGCGGCGTGGGCGTTGGTCT
GGGCGTCCCCGTCCGTCGAAGCGCAATCCAACCCGTACCAAAGAGGTCCGAACCCGACTAGATCCGCG
TTGACTGCGGACGGTCCGTTTTCCGTCGCGACTTACACTGTCTCCAGATTGTCCGTCTCCGGTTTTGG
TGGTGGTGTCATTTACTACCCGACTGGTACTTCCTTGACTTTTGGTGGTATTGCGATGTCCCCGGGTT
ACACTGCGGACGCGTCCTCCTTGGCGTGGTTGGGTAGAAGATTGGCGTCCCATGGTTTTGTCGTCTTG
GTCATTAACACTAACTCCAGATTTGACGGTCCGGACTCCAGAGCGTCCCAATTGTCCGCGGCGTTGAA
CTACTTGAGAACTTCCGACCCGTCCGCGGTCAGAGCGAGATTGGACGCGAACAGATTGGCGGTCGCGG
GTCATTCCATGGGTGGTGGTGCGACTTTGAGAATTGCGGAACAAAACCCGTCCTTGAAAGCGGCGATT
CCGTTGACTCCGTGGCATACTCAAAAAACTTTTAACACTTCCGTCCCGGTCTTGATTGTCGGTGCGGA
AGCGGACACTGTCGCGCCGGTCTCCCAACATGCGATTCCGTTTTACCAAAACTTGCCGTCCACTACTC
CGAAAGTCTACGTCGAATTGTGCAACGCGTCCCATATTGCGCCGAACTCCAACAACGCGGCGATTTCC
GTCTACACTATTTCCTGGATGAAATTGTGGGTCGACAACGACACTAGATACAGACAATTTTTGTGCAA
CGTCAACGACCCGGCGTTGTGCGACTTTAGAACTAACAACAGACATTGCCAATACCCGTACGACGTCC
CGGACTACGCGTAA
